# Mitochondrial cristae density is increased following high-intensity interval training in patients with type 2 diabetes

**DOI:** 10.1101/2025.09.29.678437

**Authors:** Martin Eisemann de Almeida, Niels Ørtenblad, Amalie Bjørnestad Platz, Maria Houborg Petersen, Kurt Højlund, Joachim Nielsen

## Abstract

**Aims/hypothesis:** Mitochondrial cristae architecture is a key determinant of oxidative capacity in skeletal muscle. While mitochondrial dysfunction is common in type 2 diabetes, it remains unclear whether cristae density is reduced and whether it can be improved by exercise training. We therefore investigated the mitochondrial cristae density in skeletal muscle of patients with type 2 diabetes compared with glucose-tolerant individuals with obesity and lean individuals, and examined the effect of high-intensity interval training (HIIT).

**Methods:** In a non-randomized intervention study, the effect of an 8-week supervised HIIT intervention combining rowing and cycling was examined in male participants (aged 40–65 years) with type 2 diabetes (*n*=15), glucose-tolerant individuals with obesity (*n*=15), and lean individuals (*n*=18). Muscle biopsies from the *m. vastus lateralis* were analyzed using transmission electron microscopy (TEM) to quantify mitochondrial cristae density (cristae surface area per mitochondrial volume) and to derive cristae surface area per muscle volume, integrating mitochondrial abundance and ultrastructure. To ensure high stereological precision, a minimum of 49 mitochondrial profiles per sample were analyzed.

**Results:** No differences in mitochondrial cristae density were observed between groups at baseline. HIIT induced an ∼7% increase in cristae density across all groups, with the most pronounced adaptations in type 2 fibers and in the intermyofibrillar compartment. At baseline, patients with type 2 diabetes exhibited lower cristae surface area per muscle volume compared with lean individuals. Notably, cristae surface area per muscle volume increased more than mitochondrial volume density alone, reflecting combined structural and volumetric remodeling.

**Conclusions/interpretation:** Skeletal muscle mitochondrial cristae density is not different between patients with type 2 diabetes and glucose-tolerant individuals with obesity and lean individuals, and the capacity for cristae remodeling in response to exercise is not affected by type 2 diabetes. These findings highlight the plasticity of mitochondrial architecture and support HIIT as a potent stimulus for improving muscle oxidative and metabolic health, also in type 2 diabetes.

**Research in Context:** *What is already known about this subject?:*

- Skeletal muscle mitochondrial cristae architecture is critical for oxidative phosphorylation and metabolic health.
- Type 2 diabetes is associated with altered mitochondrial structure, but whether cristae density is reduced remains unclear.
- Previous short-term exercise interventions have shown limited or inconsistent effects on mitochondrial cristae density, possibly due to methodological constraints.

*What is the key question?:*

- Can high-intensity interval training (HIIT) remodel skeletal muscle mitochondrial cristae in patients with type 2 diabetes, and is baseline cristae density altered in this condition?

*What are the new findings?:*

- Baseline mitochondrial cristae density does not differ between patients with type 2 diabetes and glucose-tolerant individuals with obesity and lean individuals, but cristae surface area per muscle volume is lower in type 2 diabetes.
- Eight weeks of HIIT increased mitochondrial cristae density by ∼7% across all groups, with cristae surface area per muscle volume increasing more than mitochondrial volume density alone.
- Exercise-induced cristae remodeling occurs in both muscle fiber types and subcellular compartments, demonstrating preserved structural plasticity in type 2 diabetes.

*How might this impact on clinical practice in the foreseeable future?:*

- HIIT represents a potent intervention to improve mitochondrial architecture and potentially enhance muscle oxidative capacity and metabolic health in individuals with type 2 diabetes.

## Introduction

Mitochondria are dynamic organelles essential for cellular energy conversion, metabolic regulation, and signaling [1]. They are enclosed by an outer and inner membrane with distinct protein compositions. The inner membrane forms invaginations into the matrix, termed cristae [2], encompassing the respiratory chain complexes, ATP synthase, the mitochondrial contact site and cristae organizing system (MICOS), and other factors that shape cristae architecture and function [3, 4]. Regulation of cristae architecture and density is therefore critical for oxidative phosphorylation.

In skeletal muscle from patients with type 2 diabetes, the mitochondrial network is frequently fragmented and structurally altered [5–10], which has been linked to reduced oxidative metabolism [11, 12] and impaired oxidative phosphorylation capacity in most studies [10, 13–19], although some have reported preserved function [20–25]. Whether cristae are affected remains uncertain. At the ultrastructural level, reduced mitochondrial cristae density has been observed in myotubes derived from patients with type 2 diabetes [26] and altered cristae morphology where found in mouse models of early diabetes [27], whereas muscle biopsy specimens from patients with type 2 diabetes showed no difference compared with weight-matched controls [28]. At the molecular level, proteins critical for maintaining cristae architecture, including MICOS components and the cristae-shaping GTPase optic atrophy 1 (OPA1), have been reported to be reduced in abundance in insulin-resistant muscle [29–31]. Notably, OPA1 levels correlate positively with insulin sensitivity [5, 30], suggesting a potential link between cristae organization and metabolic health, although not all studies have observed lower OPA1 in type 2 diabetes compared with weight-matched controls [5, 32].

Exercise training is a potent stimulus for mitochondrial remodeling [33, 34]. While intervention studies have generally reported no effect on mitochondrial cristae density following short-term training [28, 35], cross-sectional comparisons demonstrate ∼20% higher cristae density in endurance-trained athletes compared with untrained individuals [28], suggesting that the training stimulus in these interventions may have been insufficient. Because cristae surface area density in human skeletal muscle is typically reported within a relatively narrow range (∼25–35 µm² µm⁻³) [28, 36–38], small but physiologically meaningful differences between populations or following short-term training may be difficult to detect due to both methodological and biological variability inherent to morphometric analyses. This highlights the importance of applying high-precision morphometric approaches in intervention studies to reliably capture potential remodeling of cristae architecture.

To address this limitation, we aimed to investigate, with increased sampling precision, both baseline mitochondrial cristae density and the effect of an 8-week high-intensity interval training (HIIT) intervention in skeletal muscles of patients with type 2 diabetes, weight-matched individuals with obesity, and lean individuals. We hypothesized that patients with type 2 diabetes would display lower mitochondrial cristae density at baseline compared with both individuals with obesity and lean individuals, and that HIIT would increase cristae density across all groups.

## Methods

### Ethics

The study was approved by the Regional Committees on Health Research Ethics for Southern Denmark (S-20170142) and the Danish Data Protection Agency (17/31977) and conducted in accordance with the principles outlined in the *Declaration of Helsinki*. All participants received both oral and written information about the study, including potential risks, before providing written informed consent. Data management complied with the General Data Protection Regulation (GDPR) and was securely handled in REDCap, hosted by Odense Patient Data Explorative Network (OPEN).

### Participants

This study is a prespecified secondary analysis of a larger controlled trial, previously described in companion publications [6, 39–41]. Recruitment and intervention were conducted from January 2018 to December 2019. Fifteen patients with type 2 diabetes and overweight/obesity (BMI 27–36 kg/m^2^) were matched to 18 lean individuals (BMI 20–25 kg/m^2^) and 15 non-diabetic individuals with overweight/obesity (BMI 27–36 kg/m^2^) (Fig. 1a). Clinical and biochemical characteristics of the three groups are presented in Table 1. Eligible participants were Caucasian males aged 40–65 years who were classified as sedentary according to the International Physical Activity Questionnaire – Short Form (IPAQ–SF), reporting less than 2 h of moderate-intensity activities per week. All participants had a normal resting ECG and normal blood screening for renal, hepatic, and hematological function. Patients with type 2 diabetes were GAD65 antibody-negative, without micro- or macrovascular complications (except one with mild retinopathy), and treated with metformin (*n*=14), dipeptidyl peptidase-4 (DPP-4) inhibitors (*n*=4), or sulfonylureas (*n*=1), either as monotherapy (*n*=11) or in combination (*n*=4). They also received cholesterol-lowering (*n*=11) and antihypertensive (*n*=12) medications. The controls were glucose-tolerant based on a 2-hour 75 g OGTT, drug-naive, and without a family history of diabetes.

**Fig. 1.**
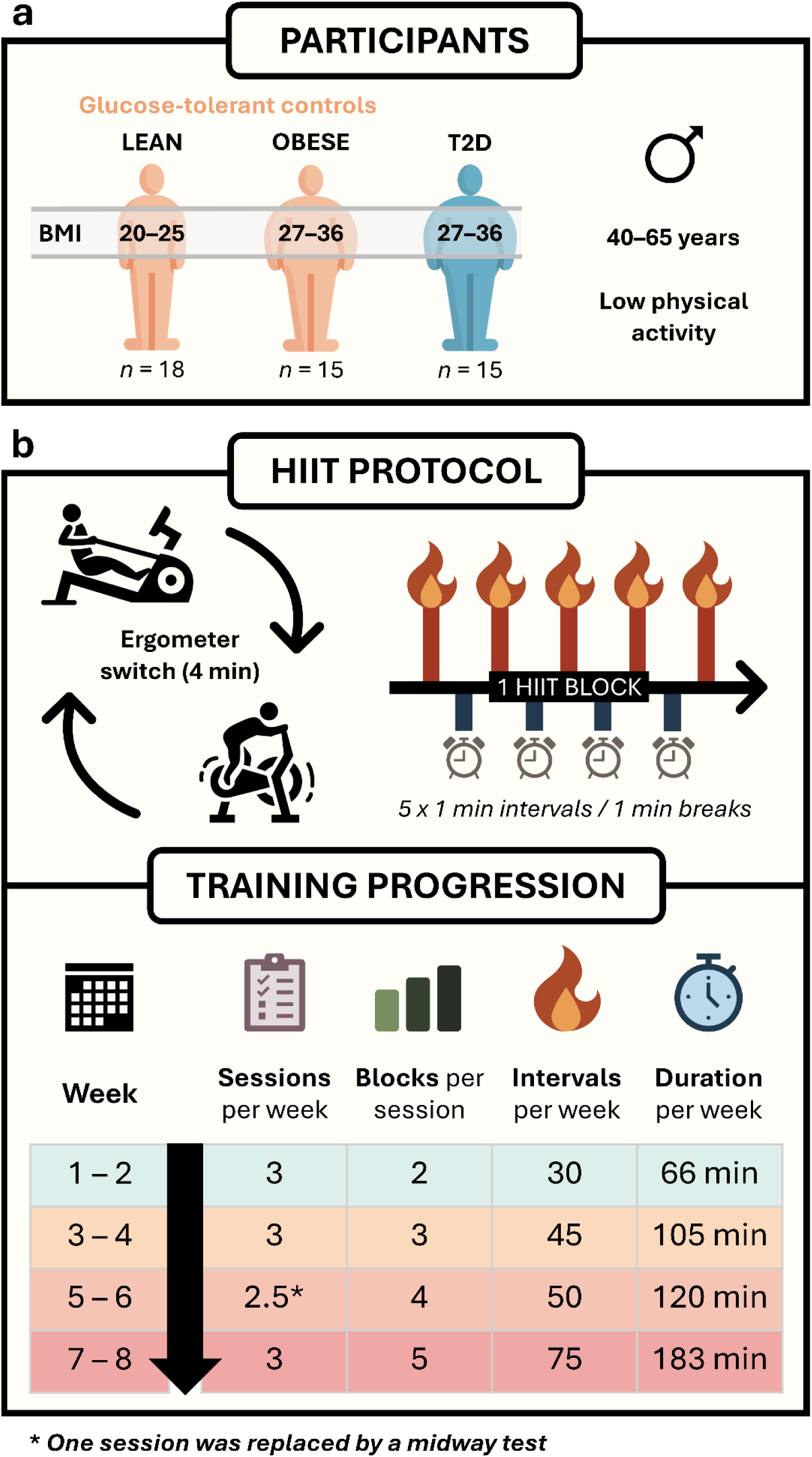
Schematic overview of participants and the 8-week high-intensity interval training (HIIT) protocol. (a) Three groups were included: lean glucose-tolerant individuals (BMI 20–25 kg/m^2^, *n* = 18), glucose-tolerant individuals with obesity (BMI 27–36 kg/m^2^, *n* = 15), and patients with type 2 diabetes (BMI 27–36 kg/m^2^, *n* = 15). All participants were male, 40–65 years old, and reported a low physical activity level. (b) Participants completed 24 planned sessions combining rowing and cycling on ergometers; one session was replaced by a mid-way test to reassess training intensity. Each session consisted of HIIT blocks of 5 × 1-min intervals separated by 1-min rest. During a 4-min transition between blocks, participants switched ergometers (rowing ↔ cycling). The number of blocks per session increased biweekly from 2 (weeks 1–2) to 5 (weeks 7–8), resulting in a weekly increase of HIIT intervals from 30 to 75 and total weekly training duration from 66 to 183 min. “Human Body” icon designed by surang, “Rowing Machine” and “Exercise Bike” icons were designed by gravisio, all via Flaticon.com.

**Table 1.** Clinical and biochemical characteristics at baseline.

|  | Lean<br>(n = 15) | Obese<br>(n = 14) | T2D<br>(n = 14) | Lean vs.<br>Obese | T2D vs.<br>Lean | T2D vs.<br>Obese |
| --- | --- | --- | --- | --- | --- | --- |
| Age (years) | 56.2 ± 6.2 | 54.0 ± 7.1 | 54.6 ± 6.2 | Main effect of group: NS |  |  |
| Duration of type 2 diabetes (years) |  |  | 4.6 ± 2.9 |  |  |  |
| Body weight (kg) | 81.2 ± 6.7 | 98.9 ± 10.7 | 104.2 ± 14.3 | *** | *** | NS |
| Body mass index (kg/m<sup>2</sup>) | 24.3 ± 1.2 | 30.6 ± 2.7 | 31.4 ± 3.1 | *** | *** | NS |
| Fat mass (kg) | 20.9 ± 4.0 | 31.4 ± 7.3 | 35.7 ± 8.3 | *** | *** | NS |
| Fat-free mass (kg) | 58.4 ± 4.5 | 65.1 ± 5.1 | 64.9 ± 6.8 | ** | ** | NS |
| VO<sub>2max</sub> (ml/kg/min) | 37.3 ± 6.4 | 33.5 ± 7.3 | 25.6 ± 3.4 | NS | *** | *** |
| Maximal fat oxidation (mg/min) | 215 ± 116 | 184 ± 128 | 145 ± 103 | Main effect of group: NS |  |  |
| HbA1c (mmol/mol) | 35.1 ± 2.8 | 34.8 ± 3.4 | 51.5 ± 11.6 | NS | *** | *** |
| Fasting plasma glucose (mmol/L) | 5.2 ± 0.4 | 5.6 ± 0.5 | 9.4 ± 2.8 | NS | *** | *** |
| Serum insulin (pmol/L) | 59 ± 32 | 71 ± 34 | 121 ± 77 | NS | ** | * |
| Plasma triglycerides (mmol/L) | 1.41 ± 0.65 | 1.48 ± 0.61 | 2.49 ± 1.59 | NS | * | * |
| LDL cholesterol (mmol/L) | 3.21 ± 0.71 | 3.63 ± 0.53 | 2.95 ± 0.64 | NS | NS | ** |
| HDL cholesterol (mmol/L) | 1.23 ± 0.34 | 1.25 ± 0.18 | 1.02 ± 0.27 | NS | * | * |
| GDR, clamp (mg/min/m<sup>2</sup>) | 368 ± 132 | 353 ± 103 | 198 ± 82 | NS | *** | *** |

Values are presented as means ± SD. Group comparisons were conducted using linear regression analysis. Statistical significance is indicated as follows: \**P* < 0.05; \*\**P* < 0.01; \*\*\**P* < 0.001; NS: not significant. T2D, type 2 diabetes; GDR, glucose disposal rate. Data adapted from [6, 39].

### Experimental protocol

Participants completed an 8-week HIIT intervention and visited the laboratory on two experimental days (≥48 h apart) both before and after the intervention. On each visit, they arrived at 07:30 AM in a fasted state (≥12 h), having abstained from alcohol and caffeine (≥24 h) and from strenuous physical activity (≥48 h). Patients with type 2 diabetes were instructed to withhold all medications (anti-diabetic, lipid-lowering, and antihypertensive) one week prior to each visit. Participants were instructed to maintain their habitual dietary intake throughout the intervention. All procedures were supervised by personnel experienced in physiological and metabolic testing.

On the first experimental day, whole-body mass and composition were assessed by dual-energy X-ray absorptiometry (DXA; Prodigy Advance, GE Healthcare). Participants then performed a graded exercise test on a cycle ergometer (SRM Ergometer System) consisting of 4-min stages starting at 40–60 W, with 20 W increments until respiratory exchange ratio (RER) reached 1.0, followed by 1-min 20 W increments to exhaustion. This test was used to determine substrate oxidation rates and cardiorespiratory fitness (V̇O_2_max; calculated as the highest 30-s average). Maximal effort was considered valid if at least two of the following criteria were met: V̇O₂ plateau <2.1 mL·kg⁻¹·min⁻¹, RER >1.1, or post-exercise lactate >8 mmol/L). Two hours before testing, participants consumed a standardized mixed breakfast (6 kcal/kg body mass; 64% carbohydrate, 22% fat, 14% protein). Full methodological details on the graded exercise test and calculation of maximal fat oxidation, including previously reported training-induced changes in V̇O₂max and substrate oxidation rates, have been described elsewhere [6, 39]. After the intervention, this first day was conducted ∼60 h after the final HIIT session.

On the second experimental day, a resting skeletal muscle biopsy was obtained, followed by a 3-h hyperinsulinemic-euglycemic clamp to assess insulin sensitivity. Clamp methodology, isotope infusion protocols, and biochemical analyses of plasma glucose, HbA1c, C-peptides, serum insulin, and lipids have been detailed in a separate prior publication [39]. Post-intervention, this test was scheduled 48 h after the first day (including V̇O₂max test) and 5 days after the final HIIT session.

### Training intervention

The 8-week intervention (Fig. 1b) comprised of three weekly HIIT sessions combining rowing (Concept 2 Model E) and cycling (Wattbike Pro/Trainer) ergometry, totaling 24 sessions, including one mid-way test to reassess training intensity. Participants across all groups trained together in groups of up to 10, with the majority of sessions (>95%) conducted in the afternoon. Each session began with a 10-minute warm-up, followed by HIIT blocks consisting of 5 x 1-minute intervals interspersed with 1-min active recovery. A 4-minute rest period was provided between blocks to switch between rowing and cycling, with the starting modality alternating between sessions. Training volume increased biweekly, from two blocks per session in weeks 1-2 to five blocks in weeks 7-8. Cycling intervals were performed at 100-110% of maximal cycling capacity (MCC), with similar heart rates achieved during rowing and cycling (86-88% of max). Rowing ergometers were calibrated with a drag factor of 105-110, and participants received real-time feedback to ensure supramaximal effort. Motivation was supported by group dynamics, supervisor encouragement, music, and live heart rate monitoring (Polar H7/Team). Detailed training data are reported in [6].

### Muscle biopsies

All skeletal muscle biopsies were collected from the *m. vastus lateralis* by the same experienced physician to minimize variation in sampling depth and location. Following local anesthesia (5 mL of 2% lidocaine), a small incision was made through the skin and fascia, and the biopsy was obtained using a modified Bergström needle with suction. Samples were immediately cleared of blood and connective tissue, then divided for various analyses, including a ∼1 mm^3^ portion prepared for TEM.

Due to a fixation error, pretraining biopsies were unavailable for five participants (three lean individuals, one individual with obesity, and one patient with type 2 diabetes), who were excluded from the analyses. In addition, four participants did not complete the intervention: one lean individual and two patients with type 2 diabetes withdrew due to time constraints, and one lean individual did not initiate training due to a knee injury. However, their baseline characteristics were retained in the present study. To increase tissue availability for mitochondrial analyses, extra pretraining biopsies were collected from 11 lean individuals, 12 individuals with obesity, and 12 patients with type 2 diabetes as part of a separate acute exercise study [40] conducted before the training intervention. These samples were obtained from the same participants and served to increase biopsy material per individual. The final analytical sample comprised: patients with type 2 diabetes (Pre: *n*=14; Post: *n*=12), individuals with obesity (Pre/Post: *n*=14), and lean individuals (Pre: *n*=15; Post: *n*=13). The baseline characteristics reported in Table 1 are based on available participants with valid pretraining biopsies.

### Transmission electron microscopy

Muscle biopsy specimens were prepared for TEM as described previously [42]. Briefly, samples were fixed in 2.5% glutaraldehyde in a 0.1 M sodium cacodylate buffer (pH 7.3) for 24 hours at 5 °C and subsequently washed four times in the same buffer. Post-fixation was carried out in a solution of 1% osmium tetroxide (OsO_4_) and 1.5% potassium ferrocyanide [K_4_Fe(CN)_6_] in 0.1 M sodium cacodylate buffer for 120 minutes at 4 °C. Following post-fixation, the specimens were rinsed twice in double-distilled water, dehydrated through a graded ethanol series, and infiltrated with graded mixtures of propylene oxide and Epon at room temperature. The following day, samples were embedded in 100% Epon and polymerized at 60 °C for 48 hours. Ultra-thin sections (60 nm) were cut using a Leica Ultracut UC7 ultramicrotome at three depths separated by 150 µm to maximize fiber sampling. Sections were contrasted with uranyl acetate and lead citrate before examination in a pre-calibrated JEM-1400plus transmission electron microscope (JEOL) equipped with a CCD camera (Quemesa, Olympus). Electron micrographs were acquired from 10 longitudinally oriented muscle fibers per biopsy. For each fiber, 24 images were captured at 6,000× magnification in a randomized systematic order: 12 from the subsarcolemmal region, and 6 from the superficial and central region of the myofibrillar space (Fig. 2b).

**Fig. 2.**
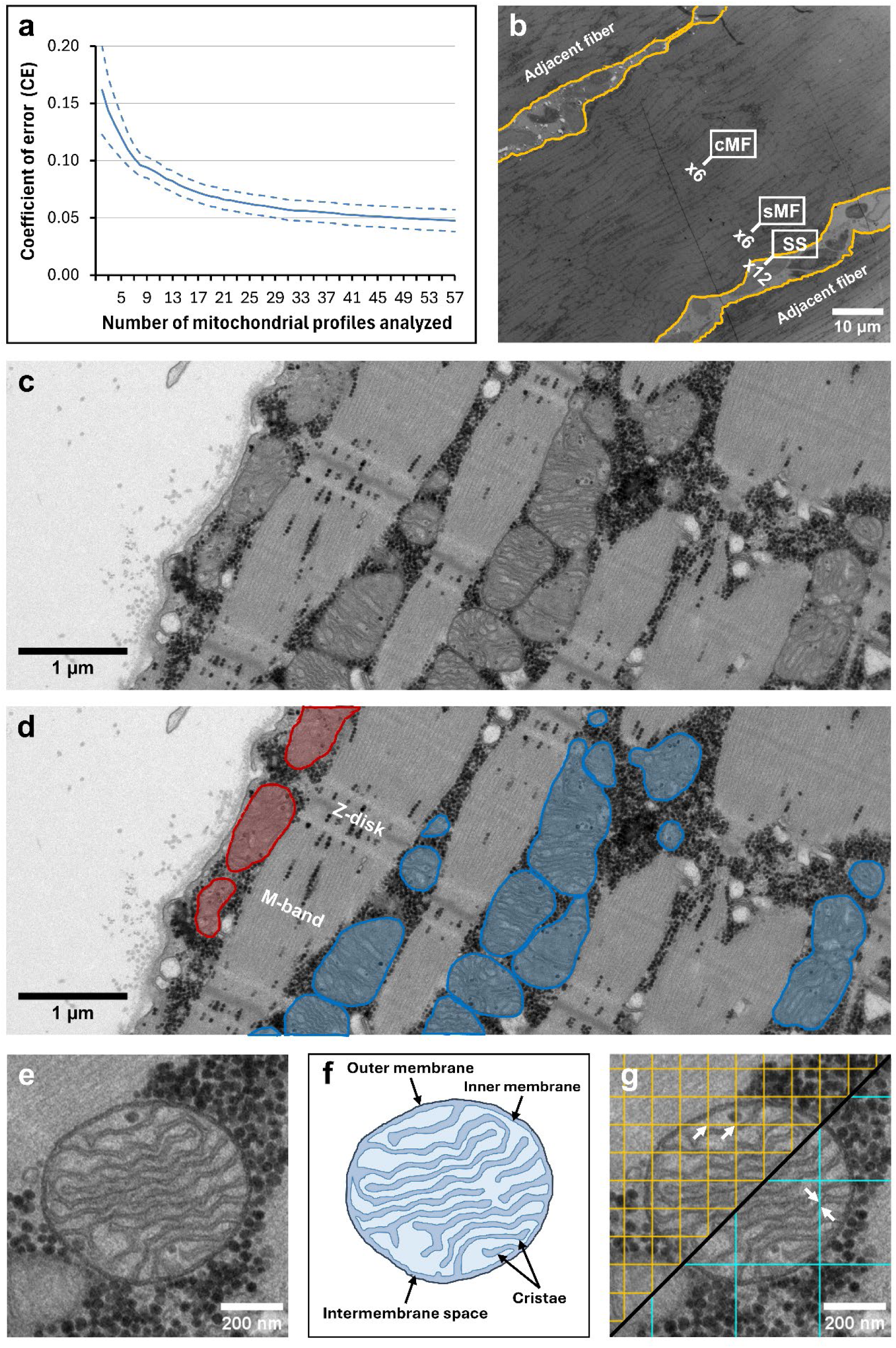
Transmission electron microscopy (TEM) and stereological assessment of skeletal muscle mitochondria. (a) To ensure high precision of cristae density estimates, ≥ 49 mitochondrial profiles per sample were analyzed, yielding an estimated coefficient of error (_est_CE) < 5%. (b). Micrographs were acquired from 10 longitudinally oriented fibers per biopsy, with 24 images per fiber: 12 subsarcolemmal (SS), and 6 from superficial (sMF) and central (cMF) myofibrillar regions. (c) Representative TEM image of a skeletal muscle fiber. (d) The same image with subsarcolemmal mitochondria highlighted in red and intermyofibrillar mitochondria highlighted in blue. (e) Example of a mitochondrion of acceptable quality, showing clearly visible cristae structure with minimal or no missing traces. (f) Schematic illustration of mitochondrial morphology, including outer membrane, inner membrane with cristae, and intermembrane space. (g) Grids were overlaid to quantify intersections of inner membrane folds (teal lines) for cristae surface area and mitochondrial hits (orange lines) for volume; the ratio of cristae surface area to mitochondrial volume was used to calculate cristae density.

Muscle fibers were classified as type 1 or 2 based on their distinct Z-disk width, which correlates with myofibrillar ATPase staining patterns in light microscopy [43, 44]. Fibers with the thickest Z-disks were classified as type 1, and those with the thinnest as type 2. Three fibers of each type were selected per biopsy for point-counting analysis, while intermediate fibers were excluded. The median (IQR) Z-disk width was 83 (78–89) nm for type 1 fibers and 65 (62–69) nm for type 2 fibers.

### Mitochondrial volume density

Mitochondrial volume density for the three groups was previously determined [6], which also includes data on stereological precision and inter-investigator reproducibility. As described by Hoppeler et al. [37], total mitochondrial volume density in skeletal muscle encompasses both subsarcolemmal mitochondria, located between the sarcolemma and the outermost myofibrils, and intermyofibrillar mitochondria, which are positioned between the myofibrils, typically near the I-band and adjacent to the Z-disk (Fig. 2c–d).

Two subcellular compartments were defined for stereological estimation: (1) the intermyofibrillar space, and (2) the subsarcolemmal space. The volume density of intermyofibrillar mitochondria was expressed relative to the myofibrillar space and estimated by point-counting in both the superficial and central regions of the myofibrillar compartments. Given that muscle fibers are assumed to be cylindrical with a diameter of 80 µm [45], and that the superficial region occupies approximately three times the volume of the central region, counts from the superficial region were weighted three times more heavily. In contrast, the volume density of subsarcolemmal mitochondria was expressed relative to the surface area of the muscle fiber, based on direct measurements of fiber length in the subsarcolemmal region (i.e., visible myofibrils parallel to the sarcolemma). Fiber surface area was then calculated as the measured fiber length multiplied by the thickness of the ultrathin sections (60 nm).

Volume density (𝑉_𝑉_) was estimated using a 350 × 350 point grid (442 test points per micrograph), applying the stereological principle that volume density can be approximated from point density (𝑃_𝑃_), as 𝑉_𝑉_ = 𝐴_𝐴_ = 𝑃_𝑃_, where 𝐴_𝐴_ is the area density [46]. Since intermyofibrillar mitochondria were expressed per myofibrillar volume and subsarcolemmal mitochondria per surface area, total volume densities were computed by normalizing the subsarcolemmal volume densities to the muscle fiber volume. This was done by dividing the subsarcolemmal values by a factor of 20, based on the geometric assumption (𝑉_𝑏_ = 𝑟 · 0.5 · 𝐴) that the volume beneath (𝑉_𝑏_) 1 mm^2^ of surface area (𝐴) in a fiber with 40 µm radius (𝑟) is 20 mm^3^. All analyses were performed in iTEM (Olympus).

### Mitochondrial cristae density

Mitochondrial cristae density, defined as cristae surface area per mitochondrial volume, was quantified on the same electron micrographs used in the previous point-counting analysis. Mitochondrial profiles were included if they met quality criteria, defined by clear visibility and minimal disruption or loss of the inner mitochondrial membrane (Fig. 2e–f). In cases where only part of a mitochondrial profile fulfilled these criteria, that specific segment was included in the analysis. Initially, all mitochondrial profiles present in each micrograph were analyzed. However, due to the high image quality and the substantial time required, this approach proved impractical. Therefore, the strategy was revised to include one mitochondrial profile per micrograph. Selection was conducted systematically by visually scanning each image from the top-left to the bottom right corner and including the first profile that met the predefined quality criteria. This adjustment, implemented after the completion of the first of 14 folders, was made to minimize selection bias. In total, 10,851 (whole or partial) mitochondrial profiles were analyzed (Pre: 6,087; Post: 4,764), with a median (IQR) of 134 (72–218) and 117 (86–143) profiles per participant at Pre and Post, respectively.

Quantification of mitochondrial cristae density (SV) followed Weibel’s stereological principles [46], using the standard formula: 𝑆𝑉 = 2 · 𝐼_𝑑𝑏*l*_ · (𝑃_𝑚*i*_ · 𝑘 · 𝑑)^−1^, where 𝐼_𝑑𝑏*l*_ is the number of intersections between the inner membrane and test lines, 𝑃_𝑚*i*_ is the number of test points hitting the mitochondrial profile (Fig. 2g), and 𝑘 and 𝑑 are constants defined by the test system. Grid sizes of 270 × 270 nm and 90 × 90 nm were used to measure 𝐼_𝑑𝑏*l*_ and 𝑃_𝑚*i*_, respectively. All analyses were conducted in ImageJ (version 1.54d, National Institute of Health). Finally, to estimate the cristae surface area per muscle volume, mitochondrial cristae density was multiplied by mitochondrial volume density.

### Previous assessments of inter-mitochondrial profile variation [28, 36] have shown that a minimum of eight mitochondrial profiles per sample are necessary to obtain an estimated coefficient of error

(_est_CE) of 0.10. However, as the expected effect size of training on mitochondrial cristae density is likely below 10% [28], we aimed for an _est_CE of 0.05, corresponding to a minimum of 49 mitochondrial profiles per estimate (Fig. 2a). This stringent criterion, applied across fiber types (mixed fibers), led to the exclusion of 10 pre-biopsies and six post-biopsies across the three groups. The number of exclusions increased further when stratifying by fiber type and subcellular location. For transparency, results based on the less stringent threshold of only eight profiles (and thus fewer excluded samples) are also presented (Supplemental Fig. 1, 2). Two blinded investigators examined the eligible mitochondrial profiles in randomized order, with micrographs evenly distributed across groups, time points, and muscle fiber types. Cross-validation of their inter-investigator scoring revealed a CV of 4.7% for the cristae density estimates based on 49 mitochondrial profiles.

### Statistics

In alignment with the recommendations by Ho et al. [47], this study employed estimation statistics to evaluate training-induced effect sizes along with their associated 95% CI. Effect sizes were visualized using estimation plots, including paired comparisons across time, stratified by group, fiber type, and subcellular location. The magnitude of the effects was interpreted as small (0.20–0.49), moderate (0.50–0.79), or large (≥0.80), based on Cohen’s d [48]. Delta (Δ) changes were calculated for all participants, and delta-delta (Δ-Δ) comparisons were performed to assess differences in responses between the three groups. To complement the estimation approach and account for missing values, a linear mixed-effects model was also applied to test interactions and main effects. Participants were treated as random effects, and time, group, and fiber type as fixed effects. Normality was evaluated using the Shapiro-Wilk test, Skewness-Kurtosis test, and Q-Q plots. Homoscedasticity was assessed using residual plots. Variables with skewed distributions were transformed prior to analysis. Main effects and interactions were evaluated by the Wald χ^2^-test. Group differences in participant characteristics were assessed using linear regression. Pearson’s correlation coefficients were computed between mitochondrial markers and metabolic variables, as visual inspection indicated a linear relationship between the variables (approximated as 𝑦𝑦 = 𝑎𝑎 · 𝑥𝑥 + 𝑏). Statistical analyses were conducted using StataBE 18.5 (StataCorp LLC), while figures were created in Excel (Microsoft Corporation). Data are presented as means ± 95% CI or medians with IQR, unless otherwise stated. The threshold for statistical significance was set at *P* < 0.05.

## Results

### The effect of HIIT on mitochondrial cristae density

Training mediated an increase of 1.7 µm^2^·µm^-3^ (paired mean difference, Fig. 3a), corresponding to a ∼7% relative increase from baseline and a moderate effect size (Cohen’s d = 0.65). This change occurred independently of type 2 diabetes or obesity. The effect was primarily driven by adaptations in type 2 fibers, in which mitochondrial cristae density increased by ∼12% from baseline (Cohen’s d = 0.85; Fig. 3c), whereas type 1 fibers showed smaller changes (Cohen’s d = 0.35; Fig. 3b). Among subcellular regions, the strongest effect was seen in the intermyofibrillar compartment (Cohen’s d = 0.64; Fig. 3d), while adaptations in the subsarcolemmal region were smaller (Cohen’s d = 0.41; Fig. 3e).

**Fig. 3.**
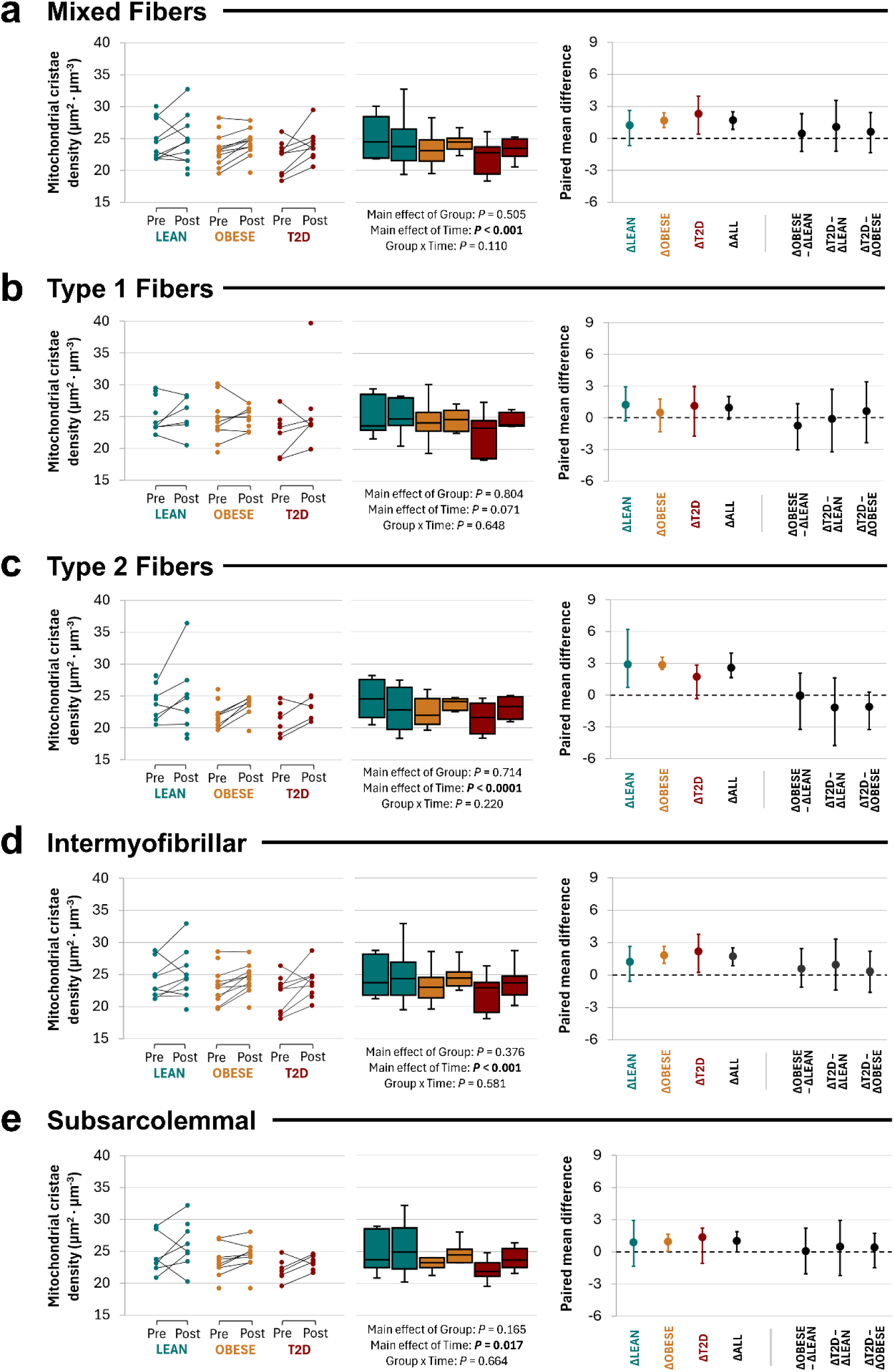
Effect of high-intensity interval training (HIIT) on mitochondrial cristae density in skeletal muscle fibers and subcellular regions. Cristae density in (a) mixed fibers, (b) type 1 fibers, (c) type 2 fibers, and (d) intermyofibrillar and (e) subsarcolemmal regions. Left panels show pre- and post-HIIT values for each participant, connected by slopegraphs. Middle panels display boxplots with medians and interquartile ranges. Main effects of group, time, and group × time interactions are reported below each panel. Right panels illustrate paired mean differences and delta–delta (ΔΔ) comparisons. Colors: Lean (teal), Obese (orange), and T2D (red). Participants were included if ≥ 49 mitochondrial profiles were analyzed (_est_CE < 5%). Individual data and boxplots: Lean: *n* = 9–11 pre, *n* = 7–12 post; Obese: *n* = 11–12 pre, *n* = 8–12 post; T2D: *n* = 7–10 pre, *n* = 6–9 post. Paired mean differences: Lean: *n* = 6–9; Obese: *n* = 6–10; T2D: *n* = 4–8.

At baseline, mitochondrial cristae density was 6% higher in type 1 fibers compared to type 2 fibers (main effect of fiber type *P* < 0.0001; Fig. 3b–c), but this difference was no longer present after HIIT. No differences in cristae density were observed between groups at either baseline or post-intervention (Fig. 3a–e), and the magnitude of change (Δ-Δ) did not differ between groups. However, based on the observed variation within the 95% CI, it cannot be excluded that individuals with type 2 diabetes or obesity may have experienced greater training-induced improvements in cristae density, potentially starting from a lower pre-training level, than lean individuals.

### The effect of HIIT on cristae surface area per muscle volume

We previously demonstrated that mitochondrial volume density is largely preserved in patients with type 2 diabetes, with no apparent differences in either the intermyofibrillar or subsarcolemmal compartments compared to individuals with obesity and lean individuals [6]. In that study, HIIT led to substantial increases in mitochondrial volume density across all three groups (∼20–40% in the intermyofibrillar and ∼30–50% in the subsarcolemmal region).

To examine whether mitochondrial remodeling extends beyond volumetric expansion, we estimated cristae surface area per muscle volume. This composite metric captures both mitochondrial abundance and internal structure. Given the increase in mitochondrial cristae density following training, the composite measure of cristae surface area per muscle volume increased more than mitochondrial volume density alone across all groups, fiber types, and subcellular compartments (All Cohen’s d > 1; Fig. 4a–e).

**Fig. 4.**
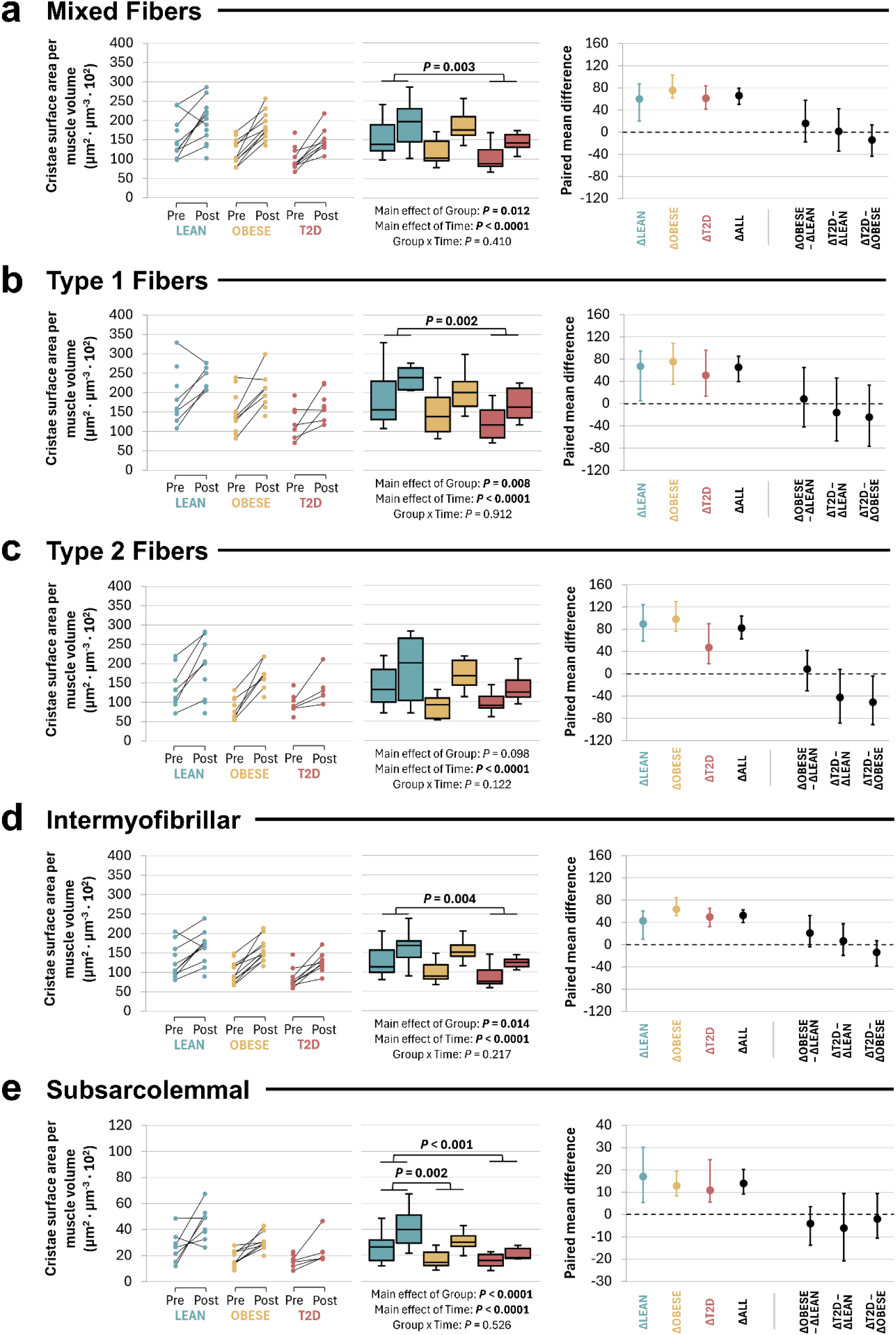
Effect of high-intensity interval training (HIIT) on mitochondrial cristae surface area per muscle volume in skeletal muscle fibers and subcellular regions. Muscle cristae surface area per muscle volume in (a) mixed fibers, (b) type 1 fibers, (c) type 2 fibers, and (d) intermyofibrillar and (e) subsarcolemmal regions. Panel layout and presentation of individual data, boxplots, paired mean differences, and ΔΔ comparisons are identical to Fig. 3. Colors: Lean (teal), Obese (yellow), and T2D (red). Participants were included if ≥49 mitochondrial profiles analyzed (_est_CE < 5%). Individual data and boxplots: Lean: *n* = 9–11 pre, *n* = 7–12 post; Obese: *n* = 11–12 pre, *n* = 8–12 post; T2D: *n* = 7–10 pre, *n* = 6–9 post. Paired mean differences: Lean: *n* = 6–9; Obese: *n* = 6–10; T2D: *n* = 4–8.

At baseline, cristae surface area per muscle volume was higher in type 1 fibers compared with type 2 fibers, and this fiber-type difference persisted after training (main effect of fiber type, both *p* < 0.001; Fig. 4b, c). While mitochondrial volume density alone did not reveal clear group differences [6], the combined measure indicated higher values in lean individuals compared to those with type 2 diabetes, particularly in type 1 fibers (Fig. 4b). This suggests that integrating volumetric and structural features may improve sensitivity to subtle group differences, although the observed pattern appeared primarily driven by changes in mitochondrial volume density.

No between-group differences in Δ-Δ (delta-delta) change of cristae surface area per muscle volume were observed for type 1 fibers (Fig. 4b) or for mitochondria in the intermyofibrillar and subsarcolemmal compartments (Fig. 4d, e). In type 2 fibers, individuals with obesity initially appeared to display a greater training-induced increase compared with patients with type 2 diabetes (Fig. 4c). However, this effect was not confirmed in supplemental analyses using a more conservative effect size threshold (_est_CE < 0.10; Supplemental Fig. 2c) and should therefore be interpreted with caution, particularly given the small sample size in the type 2 diabetes group (*n* = 4; Fig. 4c).

### Correlations between mitochondrial metrics and aerobic capacity

To increase statistical power and focus on overall associations, the three groups were combined for this analysis, and group-specific effects were not examined. Taken together, the observed training-induced changes in mitochondrial structure prompted us to explore associations both among mitochondrial measures themselves and with key measures of aerobic capacity. At baseline, mitochondrial cristae density correlated positively with the mitochondrial volume density across all fibers and subcellular regions (Fig. 5a–e). However, the HIIT-induced increases in cristae density did not correlate with the increases in mitochondrial volume density.

**Fig. 5.**
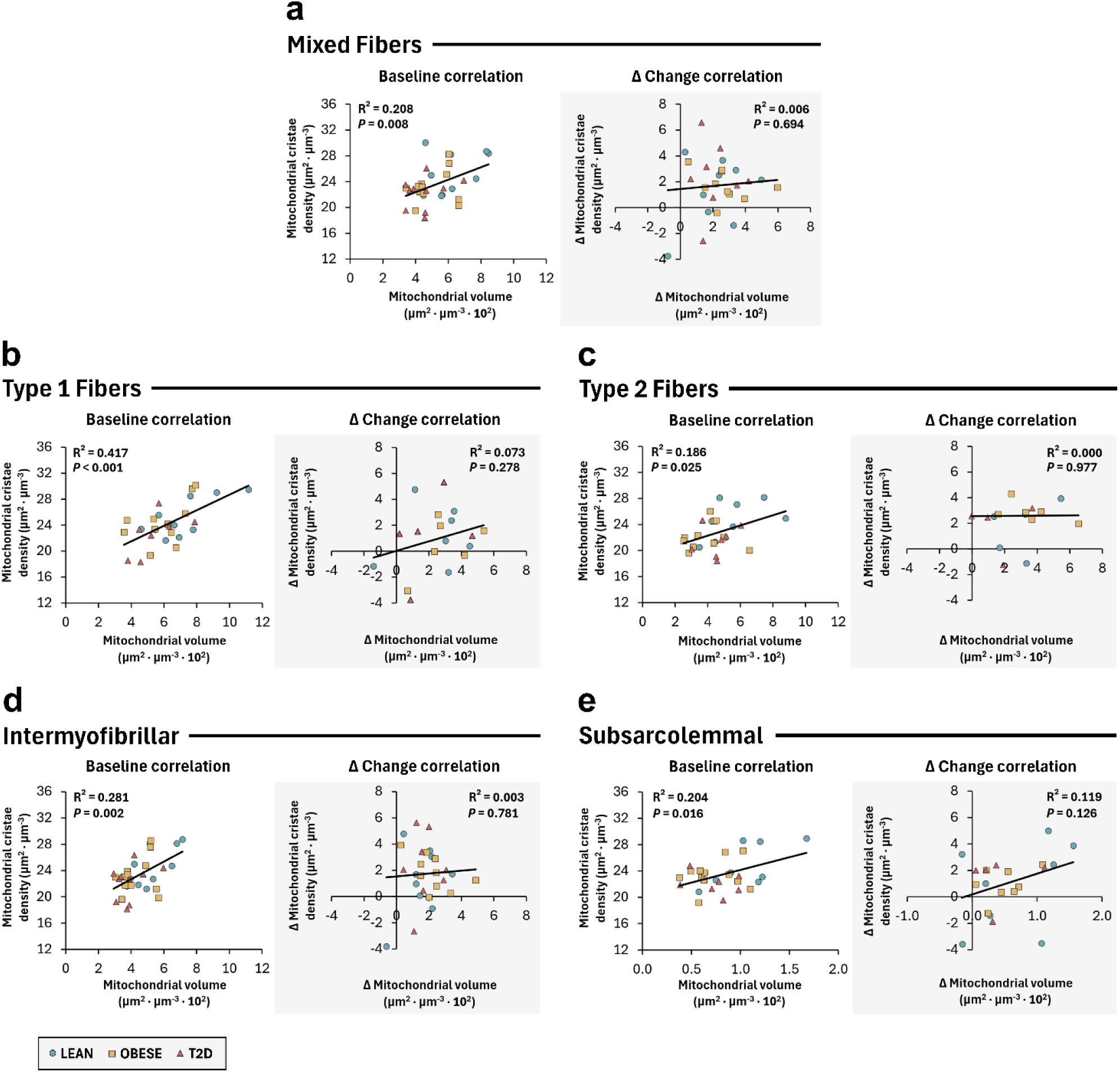
Correlations between mitochondrial volume and cristae density in skeletal muscle fibers and subcellular regions. Correlations at baseline and for delta (Δ) change in (a) mixed fibers, (b) type 1 fibers, (c) type 2 fibers, and (d) intermyofibrillar and (e) subsarcolemmal regions. Symbols indicate Lean (teal circles; baseline *n* = 9–11, Δ *n* = 6–9), Obese (yellow squares; baseline *n* = 11–12, Δ *n* = 6–10), and T2D (red triangles; baseline *n* = 7–10, Δ *n* = 4–8). Black lines represent the best linear fit including all participants. Coefficient of determination (R^2^) and *P*-values are reported in each panel.

At baseline, both maximal oxygen uptake (V̇O_2_max) and maximal fat oxidation correlated positively with mitochondrial volume density (Fig. 6a, g) and cristae surface area per muscle volume (Fig. 6c, i), but not with cristae density alone (Fig. 6b, h). When examining the shifts in mitochondrial features relative to aerobic metrics, no correlations were found for V̇O_2_max (Fig. 6d–f). However, there was a trend toward positive correlations between changes in maximal fat oxidation and changes in mitochondrial volume density (Fig. 6j) as well as changes in cristae surface area per muscle volume (Fig. 6l), while no association was observed with changes in cristae density (Fig. 6k).

**Fig. 6.**
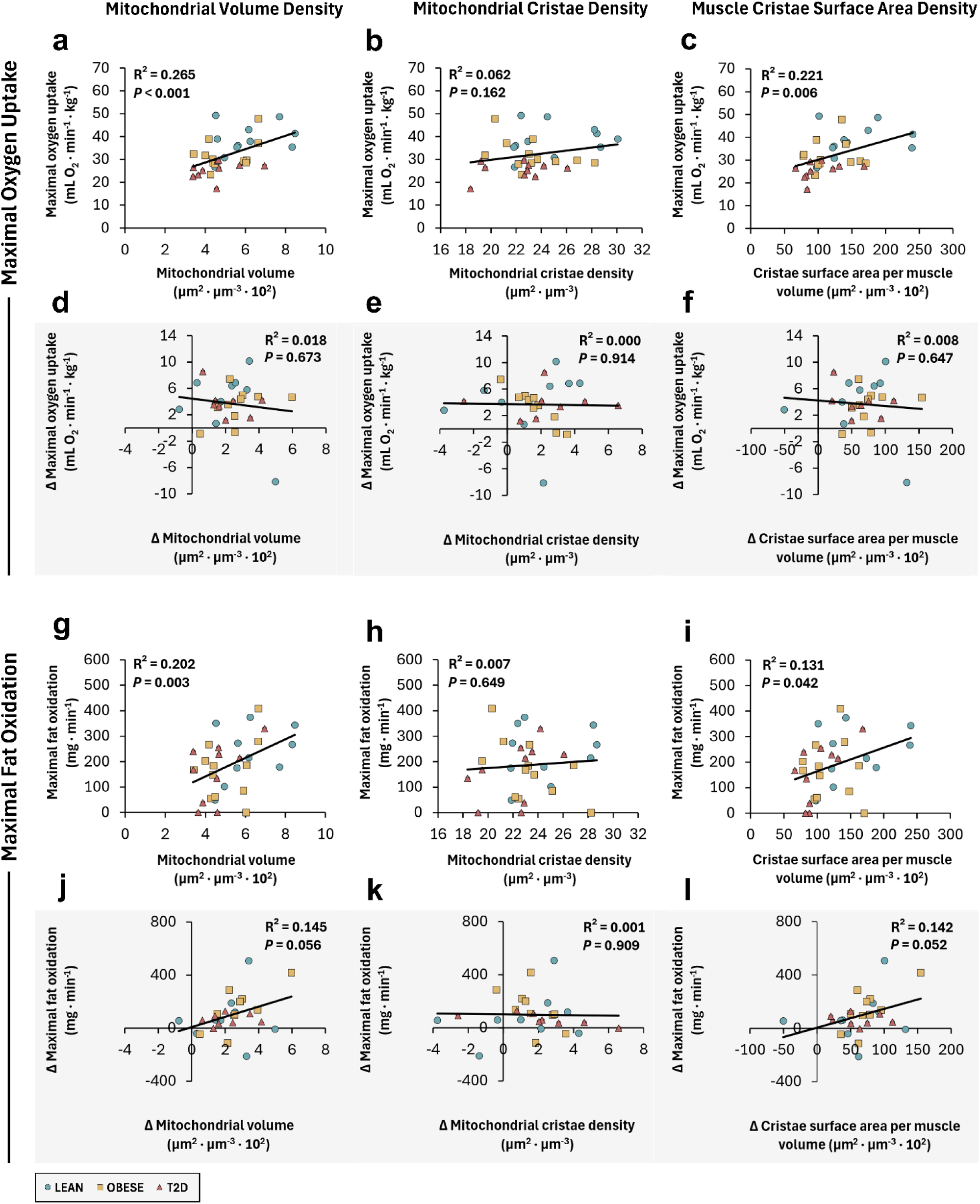
Correlations between mitochondrial morphology and maximal oxygen uptake or fat oxidation. Correlations between mitochondrial volume density, cristae density, and cristae surface area per muscle volume versus (a–f) maximal oxygen uptake (V̇O_2_max) and (g–l) maximal fat oxidation, shown at (a–c, g–i) baseline and (d–f, j–l) delta (Δ) change. Symbols indicate Lean (teal circles; baseline *n* = 11, Δ *n* = 9), Obese (yellow squares; baseline *n* = 12, Δ *n* = 10), and T2D (red triangles; baseline *n* = 10, Δ *n* = 8). Black lines represent the best linear fit including all participants. Coefficient of determination (R^2^) and *P*-values are reported in each panel.

## Discussion

Mitochondria play a central role in skeletal muscle energy metabolism, and their structural organization is a key determinant of oxidative capacity. The ultrastructure of mitochondria, particularly the invaginations of the inner membrane known as cristae, has traditionally been considered relatively invariant in skeletal muscle, regardless of diabetes status or exercise training [38, 49–52]. Here, we distinguish between mitochondrial cristae density, an ultrastructural property defined as cristae surface area per mitochondrial volume, and muscular cristae density, which additionally integrates mitochondrial volume density to reflect the total cristae surface area per muscle volume.

In this study, based on nearly 11,000 mitochondrial profiles imaged by TEM, we demonstrate that mitochondrial cristae density is responsive to HIIT and that this plasticity is preserved in skeletal muscle of patients with type 2 diabetes. Although mitochondrial cristae density did not differ between patients with type 2 diabetes, glucose-tolerant individuals with obesity, and lean individuals, cristae surface area per muscle volume was lower in patients with type 2 diabetes compared with lean individuals. Importantly, HIIT increased cristae surface area per muscle volume beyond changes in mitochondrial volume density alone, reflecting combined structural and volumetric remodeling.

Our findings of preserved mitochondrial cristae density in patients with type 2 diabetes contrasts with previous reports of reduced cristae density [26] and disturbed circadian regulation of inner membrane gene expression [30] in myotubes from type 2 diabetes donors. However, Rajab et al. [27] found dysregulated mitochondrial morphology and altered respiratory chain activity in a mouse model of early diabetes despite preserved cristae density. Similarly, we previously observed preserved mitochondrial cristae density in another cohort of patients with type 2 diabetes [28]. Collectively, these findings underscore the heterogeneity of mitochondrial alterations in diabetes and highlight the unresolved question of whether structural remodeling is impaired in this condition.

Although mitochondrial cristae density was preserved in our study, an integrated measure of cristae surface area per muscle volume revealed higher values in lean individuals compared with patients with type 2 diabetes, particularly in type 1 fibers, with no differences between patients with type 2 diabetes and individuals with obesity. This pattern appeared to be largely driven by lower mitochondrial volume density in type 2 diabetes, underscoring the importance of incorporating volumetric context to uncover subtle structural differences. In line with this, studies of diabetic myocardium exposed to hypoxia have shown that a composite index of mitochondrial volume, cristae density, and ATPase particle abundance provides a more sensitive marker of impaired ATP-generating capacity than the single ultrastructural parameters [53]. Together, these findings suggest that functional impairments in type 2 diabetes are not driven by fixed limitations of cristae packing but are better understood within a more complex structural and functional framework.

For the first time in a longitudinal study, we demonstrate that 8 weeks of HIIT increased mitochondrial cristae density by ∼7%, irrespective of type 2 diabetes or obesity. This finding contrasts with previous longitudinal human studies. Menshikova et al. [54] reported biochemical signatures of cristae expansion without ultrastructural confirmation, while Nielsen et al. [28] and Lüthi et al. [35] found no changes in cristae density after aerobic or resistance training, respectively. This discrepancy with our findings likely reflects both methodological precision and the potency of the HIIT stimulus. By quantifying a minimum of 49 mitochondrial profiles per sample (_est_CE of 0.05), we effectively doubled the resolution compared with earlier studies [28, 36], enabling detection of the modest 7 % increase in cristae density that may otherwise have remained below the threshold of detection.

An increase in mitochondrial cristae density aligns with cross-sectional studies consistently showing higher cristae densities in endurance-or strength-trained athletes compared with untrained individuals [28, 36–38, 55], with trained muscle approaching ∼35 µm² µm⁻³ compared with ∼25 µm² µm⁻³ in sedentary counterparts. Seminal animal studies have further demonstrated denser cristae packing following chronic electrical stimulation of rabbit [56] and cat muscles [57], or after prolonged running in rats [58]. Collectively, these findings reinforce the view that cristae density is a hallmark of endurance adaptation.

A key finding was that adaptations were most pronounced in type 2 fibers and in the intermyofibrillar compartment. This is relevant because type 2 fibers in type 2 diabetes are often characterized by lower oxidative enzyme activity and higher lipid content compared with lean individuals [59]. Our data show that these fibers, despite their metabolic disadvantage, retain high plasticity when exposed to sufficient exercise stimuli, narrowing the gap with type 1 fibers. At the subcellular level, the greater adaptation in the intermyofibrillar compartment is consistent with its close coupling to contractile activity and substrate utilization, and aligns with Buser et al. [60], who showed disproportionate remodeling of subsarcolemmal versus intermyofibrillar mitochondria during cold adaptation in rats, underscoring compartment-specific plasticity of cristae organization. Importantly, we observed that the increase in cristae surface area per muscle volume exceeded the change in mitochondrial volume density, suggesting that exercise training induces not only more mitochondria but also structurally more complex organelles, with potential functional implications for oxidative metabolism. Together, these data highlight that mitochondrial ultrastructure is both adaptable and functional relevant, with remodeling occurring in a fiber- and compartment-specific manner, and that this adaptive response is preserved in muscles from patients with type 2 diabetes.

Increased mitochondrial cristae density has been proposed to offset the trade-off between mitochondrial volume density and cellular space occupancy [28]. Our data demonstrate that short-term training in non-athletes with low mitochondrial volume density increases cristae density, suggesting that this adaptation can occur independently of spatial constraints. Consistently, prolonged altitude exposure has been reported to decrease cristae density while increasing mitochondrial volume density [61], confirming no fixed relationship between these parameters. High-precision studies are needed to determine how mitochondrial cristae density adapts to different training modalities and metabolic stressors.

In conclusion, skeletal muscle mitochondrial cristae density is not different between patients with type 2 diabetes and glucose-tolerant individuals with obesity and lean individuals, and the capacity for cristae remodeling in response to exercise training is preserved in type 2 diabetes. Notably, HIIT increased cristae surface area per muscle volume beyond changes in mitochondrial volume density alone, reflecting combined structural and volumetric remodeling. These findings highlight the plasticity of mitochondrial architecture and support HIIT as a potent stimulus for improving muscle oxidative potential, even in metabolically compromised skeletal muscle.

## Acknowledgments

The authors thank Medical Laboratory Technologists L. Hansen and C. B. Olsen at the Steno Diabetes Centre Odense, Odense University Hospital, for their skilled technical assistance in clinical and metabolic testing. The authors also thank Medical Laboratory Technologists A. M. Rojek and S. H. Riggelsen at the Department of Pathology, Odense University Hospital, for their excellent technical assistance with TEM.

## Data availability

The datasets generated and/or analyzed during the current study are available from the corresponding authors upon reasonable request.

## Authors’ relationships and activities

The authors declare that there are no financial or non-financial relationships, activities, or affiliations that could be perceived as influencing, or potentially biasing, the content of this work.

## Contribution statement

M.E.d.A., M.H.P., K.H., and N.Ø conceptualized and designed of the study. M.H.P. recruited the participants and conducted the metabolic testing. M.E.d.A. supervised the training sessions and performed the V̇O_2_max assessments and DXA scans. M.E.d.A. acquired the TEM micrographs, and along with A.B.P., conducted the morphological analysis of mitochondrial cristae. M.E.d.A. A.B.P., J.N., and N.Ø. analyzed and interpreted the data. M.E.d.A. prepared the figures, wrote the initial draft, and finalized the manuscript. J.N. and N.Ø. provided supervision and contributed to manuscript drafts. All authors critically revised the manuscript for intellectual content, approved the final version, and agree to be accountable for all aspects of the work.

## Abbreviations

HIIT: High-intensity interval training
TEM: Transmission electron microscopy

