## Supplemental Figures for "Mitochondrial cristae density is increased following high-intensity interval training in patients with type 2 diabetes"

**Supplemental Fig. 1. Effect of high-intensity interval training (HIIT) on mitochondrial cristae density ($_{est}$CE < 10%) in skeletal muscle fibers and subcellular regions**

Cristae density in (a) mixed fibers, (b) type 1 fibers, (c) type 2 fibers, and (d) intermyofibrillar and (e) subsarcolemmal regions. Left panels show pre- and post-HIIT values for each participant, connected by slopegraphs. Middle panels display boxplots with medians and interquartile ranges. Main effects of group, time, and group × time interactions are reported below each panel. Right panels illustrate paired mean differences and delta–delta (ΔΔ) comparisons. Colors: Lean (teal), Obese (orange), and T2D (red). Participants were included if ≥ 8 mitochondrial profiles were analyzed ($_{est}$CE < 10%). Individual data and boxplots: Lean: $n$ = 11–13 pre, $n$ = 10–12 post; Obese: $n$ = 13–14 pre, $n$ = 12–14 post; T2D: $n$ = 10–11 pre, $n$ = 9–11 post. Paired mean differences: Lean: $n$ = 9–11; Obese: $n$ = 11–14; T2D: $n$ = 8–10.

**Supplemental Fig. 2. Effect of high-intensity interval training (HIIT) on mitochondrial cristae surface area per muscle volume ($_{est}$CE < 10%) in skeletal muscle fibers and subcellular regions**

Muscle cristae surface area per muscle volume in (a) mixed fibers, (b) type 1 fibers, (c) type 2 fibers, and (d) intermyofibrillar and (e) subsarcolemmal regions. Panel layout and presentation of individual data, boxplots, paired mean differences, and ΔΔ comparisons are identical to Supplemental Fig. 1. Colors: Lean (teal), Obese (yellow), and T2D (red). Participants were included if ≥8 mitochondrial profiles analyzed ($_{est}$CE < 10%). Individual data and boxplots: Lean: $n$ = 11–13 pre, $n$ = 10–12 post; Obese: $n$ = 13–14 pre, $n$ = 12–14 post; T2D: $n$ = 10–11 pre, $n$ = 9–11 post. Paired mean differences: Lean: $n$ = 9–11; Obese: $n$ = 11–14; T2D: $n$ = 8–10.

#### Supplemental Fig. 1

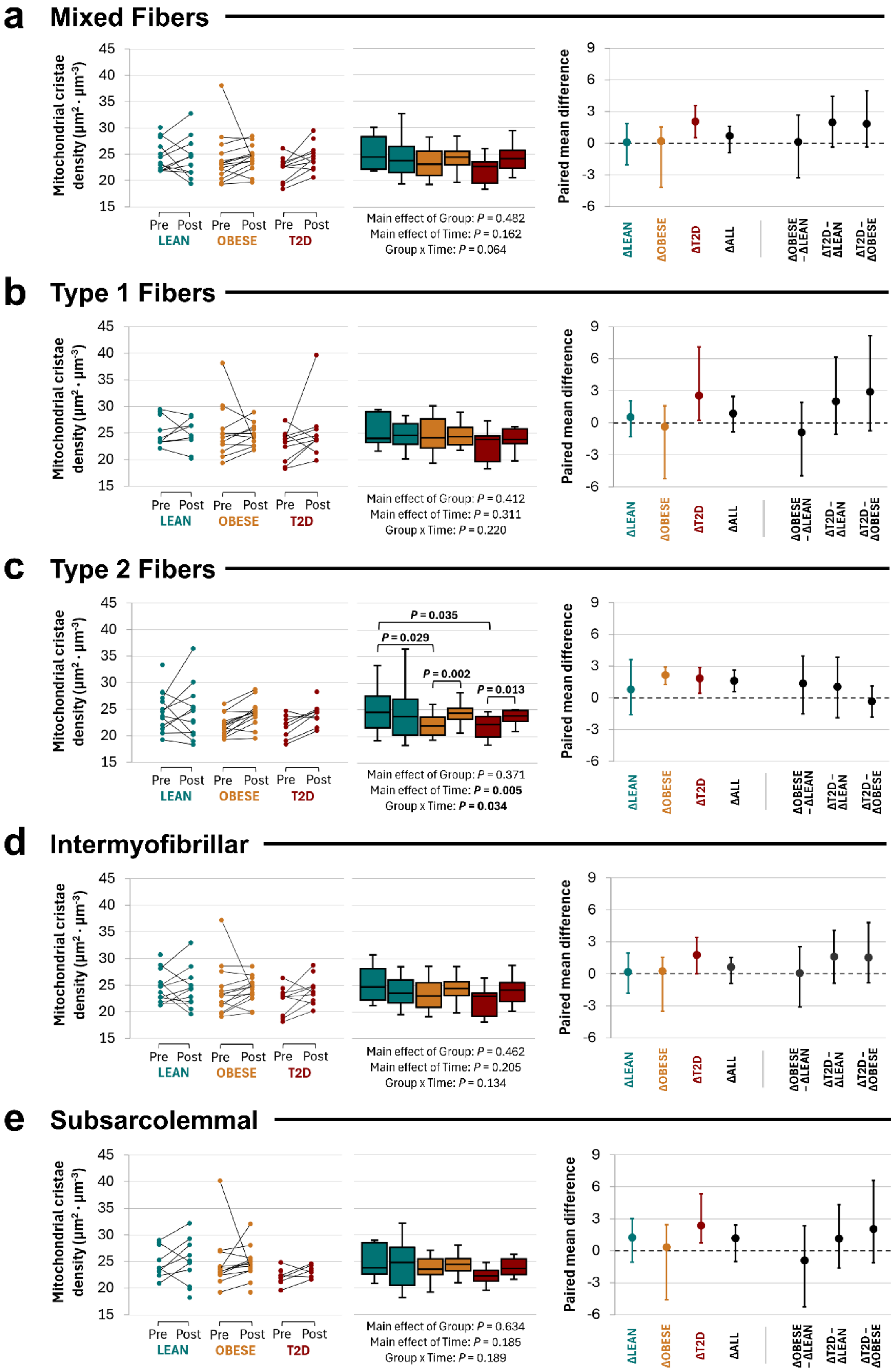

### Supplemental Fig. 2

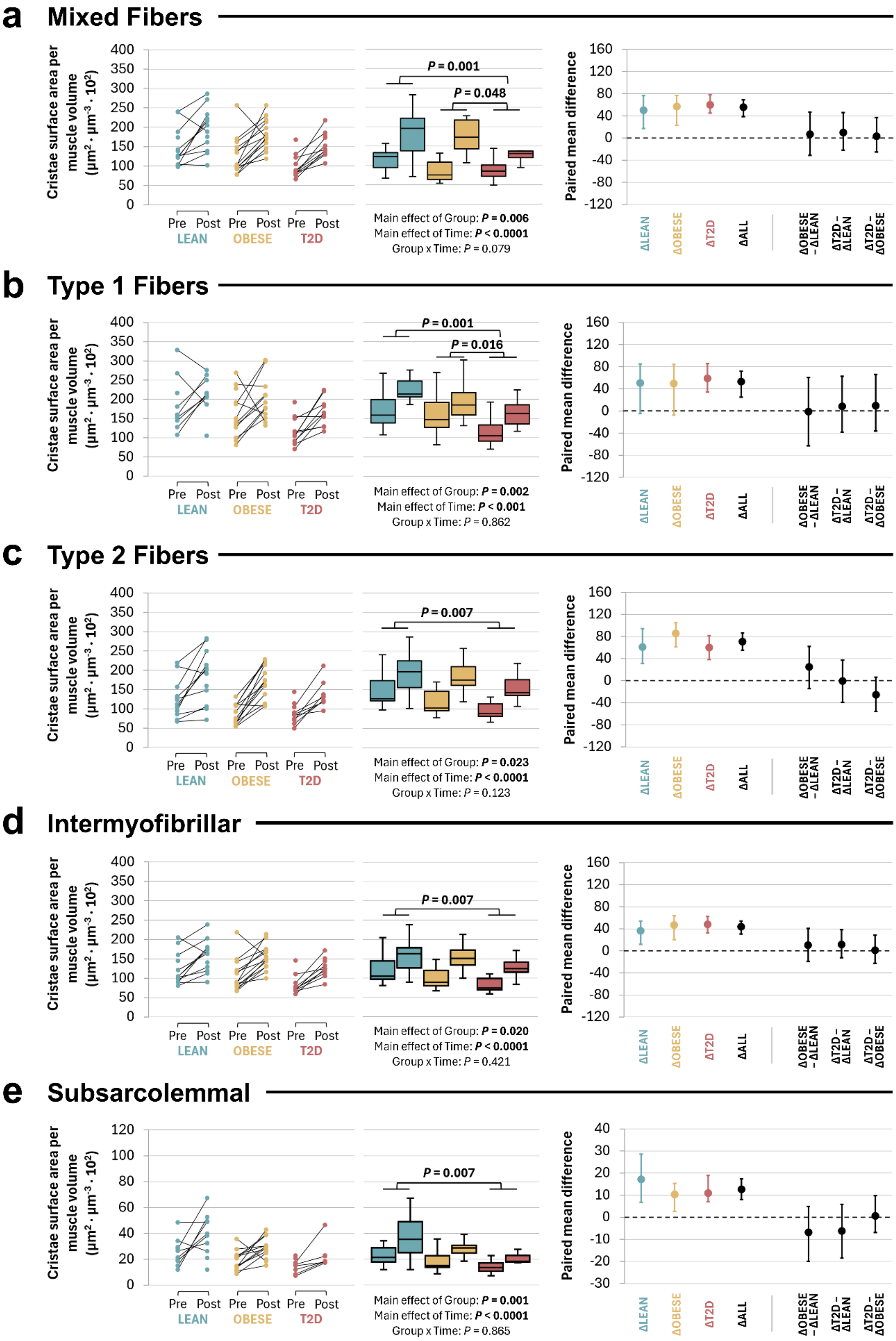
